# Diversity of Antibiotic Resistance genes and Transfer Elements-Quantitative Monitoring (DARTE-QM): a method for detection of antimicrobial resistance in environmental samples

**DOI:** 10.1101/2021.08.06.455440

**Authors:** Schuyler D. Smith, Jinlyung Choi, Nicole Ricker, Fan Yang, Shannon Hinsa-Leasure, Michelle Soupir, Heather Allen, Adina Howe

**Affiliations:** Bioinformatics and Computational Biology Program, Iowa State University, Ames, IA; Department of Agricultural and Biosystems Engineering, Iowa State University, Ames, IA; Food Safety and Enteric Pathogens Research Unit, ARS-USDA National Animal Disease Center, Ames, IA; Department of Pathobiology, Ontario Veterinary College, University of Guelph, Guelph, ON, Canada; Department of Biology, Grinnell College, Grinnell, IA

**Author notes:** Correspondence to Adina Howe.

## Abstract

Effective monitoring of antibiotic resistance genes and their dissemination in environmental ecosystems has been hindered by the cost and efficiency of methods available for the task. We developed a method entitled the Diversity of Antibiotic Resistance genes and Transfer Elements-Quantitative Monitoring (DARTE-QM), a system implementing high-throughput sequencing to simultaneously sequence thousands of antibiotic resistant genes representing a full-spectrum of antibiotic resistance classes commonly seen in environmental systems. In this study, we demonstrated DARTE-QM by screening 662 antibiotic resistance genes within environmental samples originated from manure, soil, and animal feces, in addition to a mock-community used as a control to test performance. DARTE-QM offers a new approach to studying antibiotic resistance in environmental microbiomes, showing advantages in efficiency and the ability to scale for many samples. This method provides a means of data acquisition that will alleviate the obstacles that many researchers in this area currently face.

## INTRODUCTION

The global spread of organisms possessing antimicrobial resistance (AMR), and their associated antibiotic resistant genes (ARGs), is posing an increasing threat to the health of both humans and animals alike^1–3^. Characterization of the presence and abundance of ARGs, i.e. the resistome, in environmental microbiome samples has stood as a major challenge for researchers monitoring these events^4^. Such studies have been impeded by the broad diversity of the genes, their low presence in most natural environments, the difficulty of extracting DNA from microbes in those environments, and their association with mobile genetic elements accounting for approximately one-quarter of the genetic material in these microbiomes^5^.

The genetic diversity of ARGs has made targeted sequencing approaches non-trivial and has led to the application of whole-genome shotgun metagenomic methods for the characterization resistomes^6^. This approach is dependent on the availability of a gene reference database to classify reads as ARGs sequences but does not require *a priori* knowledge of which genes constitute the resistome being investigated^7^. Despite being effective for the task, the cost per sample of employing metagenomic methods to elucidate resistomes often inhibits studies from scaling. Shotgun sequencing must indiscriminately sequence a genome, and often the resistome comprises only a fraction of a percent of the entire metagenome. Therefore, it is often the case that only a minute subset of the sequencing-reads produced through this method will be informative to resistomes, and ARGs are either underrepresented or undetected^8^, as sufficient sequencing depth and coverage is difficult to achieve.

In the effort to find more efficient means for sequencing ARGs, a method of implementing bait-and-capture system to identify ARG targets has been developed^9^. This approach uses streptavidin-coated magnetic beads to capture 80-mer bait sequences to target genes of interest. The bait-and-capture method has been well-suited for the characterization of low- and high-abundance ARGs and has demonstrated the ability to differentiate resistomes from different sample sources^10^. Another method of targeted gene sequencing used for ARG characterization involves custom primers for performing a PCR-based amplicon library preparation. This type of sequencing is used extensively in microbiome studies for community profiling via bacterial 16S rRNA genes and combines barcoded adapters to differentiate hundreds of samples pooled in a single library preparation^11^. It has previously been limited in the number of primers that could be incorporated for a single library, but a more recent version of amplicon library preparation for multiplexed primers now exists and has been implemented for biomarker detection in clinical studies^12–15^.

Our study demonstrates the first usage of this multiplexed amplicon library preparation for the detection of ARGs in environmental samples. We have termed our method of implementing this technology <u>D</u>iversity of <u>A</u>ntibiotic <u>R</u>esistance genes and <u>T</u>ransfer <u>E</u>lements-<u>Q</u>uantitative <u>M</u>onitoring (DARTE-QM). Our study was designed to demonstrate that DARTE-QM offers practical application to ARG screening through its ability to simultaneously detect and quantify hundreds of ARGs residing in samples from various environments and that it can achieve high accuracy and sensitivity identifying ARG targets.

## RESULTS

### Design of primers and samples

DARTE-QM employed 796 primer pairs designed to target 67 antibiotic resistant families and 662 ARGs, as well as a synthetic oligonucleotide reference sequence and the V4 region of the 16S rRNA gene, in a multiplexed amplicon library preparation (Supp. Table 1, Supp. Table 2). Subsequent paired-end sequencing of 150 base pair reads was conducted using the Illumina MiSeq platform (USDA, Ames, IA). To evaluate the results of DARTE-QM against a reference, we constructed a mock-community microbiome comprised of DNA extracted from 20 isolates (Supp. Table 3) with completed genome sequences (Dataset 1). For each of the mock-community libraries, we included varying concentrations of a synthetic oligonucleotide reference sequence to evaluate accuracy of quantification. We also examined how DARTE-QM was able to characterize true environmental resistomes associated with manure, swine fecal, and agricultural soil samples (Supp. Table 4).

### Evaluation of DARTE-QM sequencing products

The sequencing data produced via DARTE-QM is unique in its high level of heterogeneity, as compared to traditional amplicon data generated from a singular DNA-primer (e.g., 16S SSU rRNA). Given numerous and diverse gene targets in the sequencing library, processing of DARTE-QM data required amendment of the traditional microbiome analysis pipelines (Figure 1). After quality control and processing, 16 of the 18 samples from the mock-community were retained for downstream analysis (2 samples removed for less than 5,000 reads passing quality filters). Quality filtering also resulted in the removal of 38 of the 61 environmental samples due to sequencing coverage below 5000 reads, likely caused by PCR inhibitors common of manure and soil samples ^16–18^, leaving 39 samples in total to be used in the evaluation of DARTE-QM. The 16 mock-community samples yielded a mean of 192,415 reads per sample, and a mean of 44,440 reads able to be aligned to ARG references (Supp. Table 5). In our environmental samples, across all sources, we observed a mean of 170,775 reads and a mean of 19,138 reads aligned to ARG references per sample.

**Figure 1.**
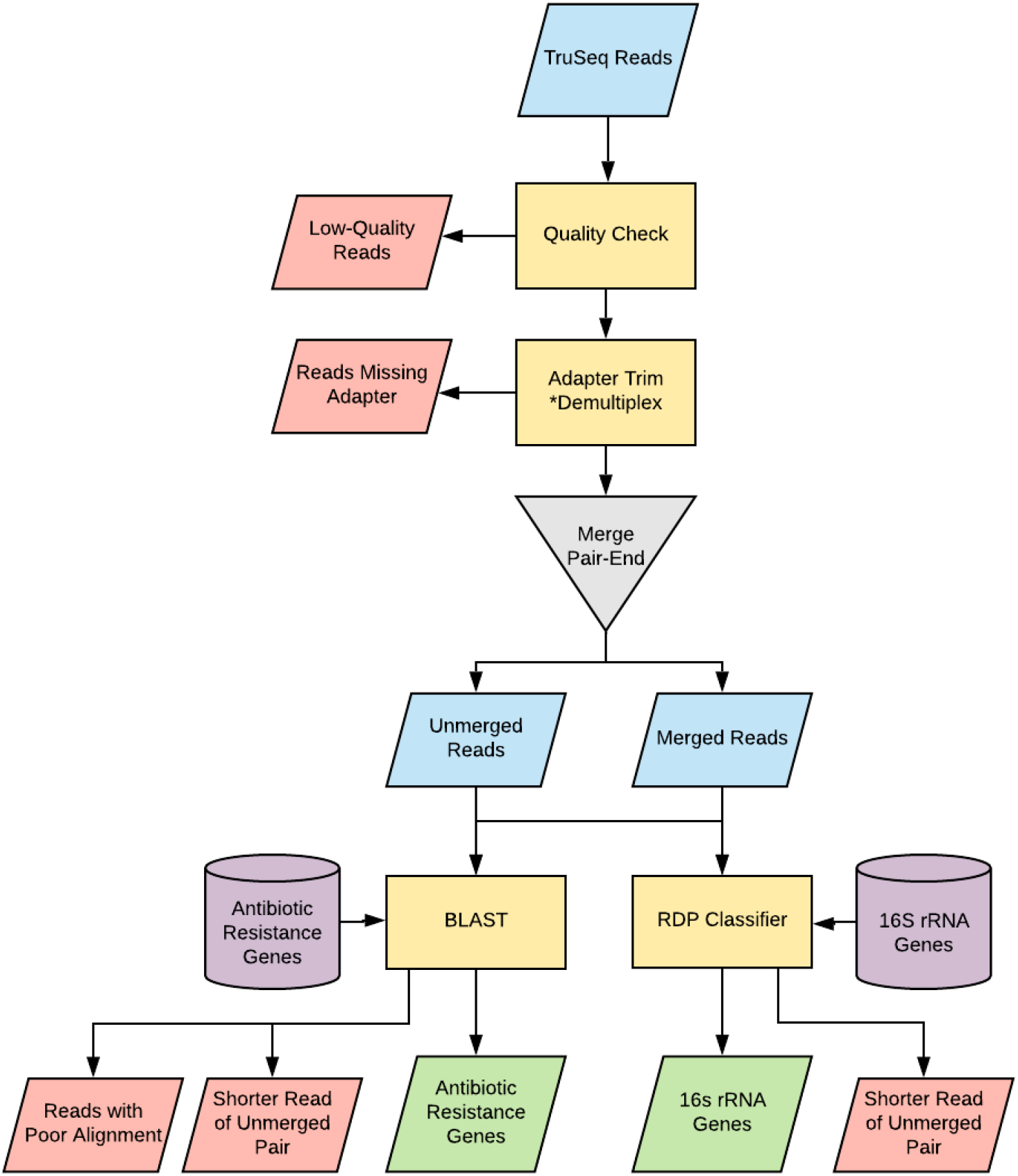
Summary of DARTE-QM read processing pipeline. Data-boxes color blued represent reads kept within the pipeline, red boxes were discarded reads, and green are the finalized reads for analysis. Reads were filtered by quality-score and demultiplexed by the presence of primer sequences. To classify ARGs, both merged and unmerged reads were required to align to known genes in ResFam and CARD ARG reference databases. In the case of unmerged reads, if both the forward and the reverse read aligned to the same target, the shorter alignment from the pair was discarded.

### DARTE-QM successfully amplified targeted genes with high accuracy and sensitivity

Reads from each sample were demultiplexed by primer, and each read was subsequently classified as either true positive (TP), false positive (FP), false negative (FN), or true negative (TN). This classification was based on alignment to the ARG reference database, where reads were deemed to be TP when both the intended primer target and read sequence aligned to the same gene; FP when the primer target and the read sequence did not agree; FN when no primer was found but the read sequence was able to be aligned to a reference ARG; and TN reads were assigned as all reads within a sample assigned as TP outside of the primer in question, i.e., all reads that were correctly identified as not being the targeted read.

Success for DARTE-QM was evaluated on three metrics: sensitivity (TP/[TP + FN]), specificity (TN/[TN + FP]), and accuracy ([TP + TN]/[TP + FN + TN + FP]) for each gene target (i.e., primer) and each sample (Supp. Table 6). From the 662 ARGs targeted by DARTE-QM, 235 (∼35%) were identified in our samples. The mean sensitivity for all primers was found to be 99.6%. The mean specificity and mean accuracy were found to be > 99.9% and 99.6%, respectively, suggesting that the primers in DARTE-QM were successful in amplifying their intended target genes.

We also observed a substantial number of reads in our sequencing libraries that had primers located on the 5’ end of the sequences but were unable to be aligned to any of our reference ARGs nor any position in the mock-community genomes. Inspection of a subset of these reads found that they contained repeated poly-A and poly-T elements. These reads were observed as unique sequences within the dataset, implying little or no biological pattern. These artifacts accounted for 47% of all reads in samples which passed quality controls. However, in samples that failed to pass quality filters, these sequences accounted for 85% of reads. Sample source appeared to be a significant, yet likely confounded, factor in the production of these artifacts. Sequencing of samples from the mock-community had significantly lower counts for artifact reads as compared to environmental samples (soil-A, p = 0.038; manure-A + soil-A, p = 0.035; swine fecal, p < 0.001, pairwise-Wilcoxon). Across all samples, an inverse linear relationship (R^2^ = 0.68) (Supp. Figure 1) was observed between the number of reads which had a primer identified and the percentage of those reads that were artifacts.

### DARTE-QM was able to consistently identify presence and distribution of ARGs

Construction of the mock-community from DNA sourced from fully-sequenced genomes allowed for comparison of a theoretical profile to our experimental observations of ARGs in these samples. In the combined genomes of the mock-community, ARGs comprised 0.03% (56 ARG targets) of the total genome by base pair count. DARTE-QM was able to produce 55 of those 56 ARGs found in the mock-community reference genomes, consistently identifying them across all 16 samples (Figure 2). Particular resistance families that were not successfully captured by DARTE-QM included those associated with the *acrA* subunit of multidrug efflux pump systems, as well as genes encoding for chloramphenicol resistance (e.g., *catA*). While overall, target relative abundances were observed to be similar compared to theoretical, the quantification of particular ARGs, such as transposon-associated *lnuC* conferring resistance to lincomycin, were found in higher abundance by DARTE-QM, as others such as *mecA* conferring methicillin-resistance, were found to be underrepresented. With regard to the synthetic oligonucleotide, there was a strong correlation observed reads (Supp Figure 2, R^*2*^ *=* 0.91) between the read abundance produced by DARTE-QM and the experimental concentration.

**Figure 2.**
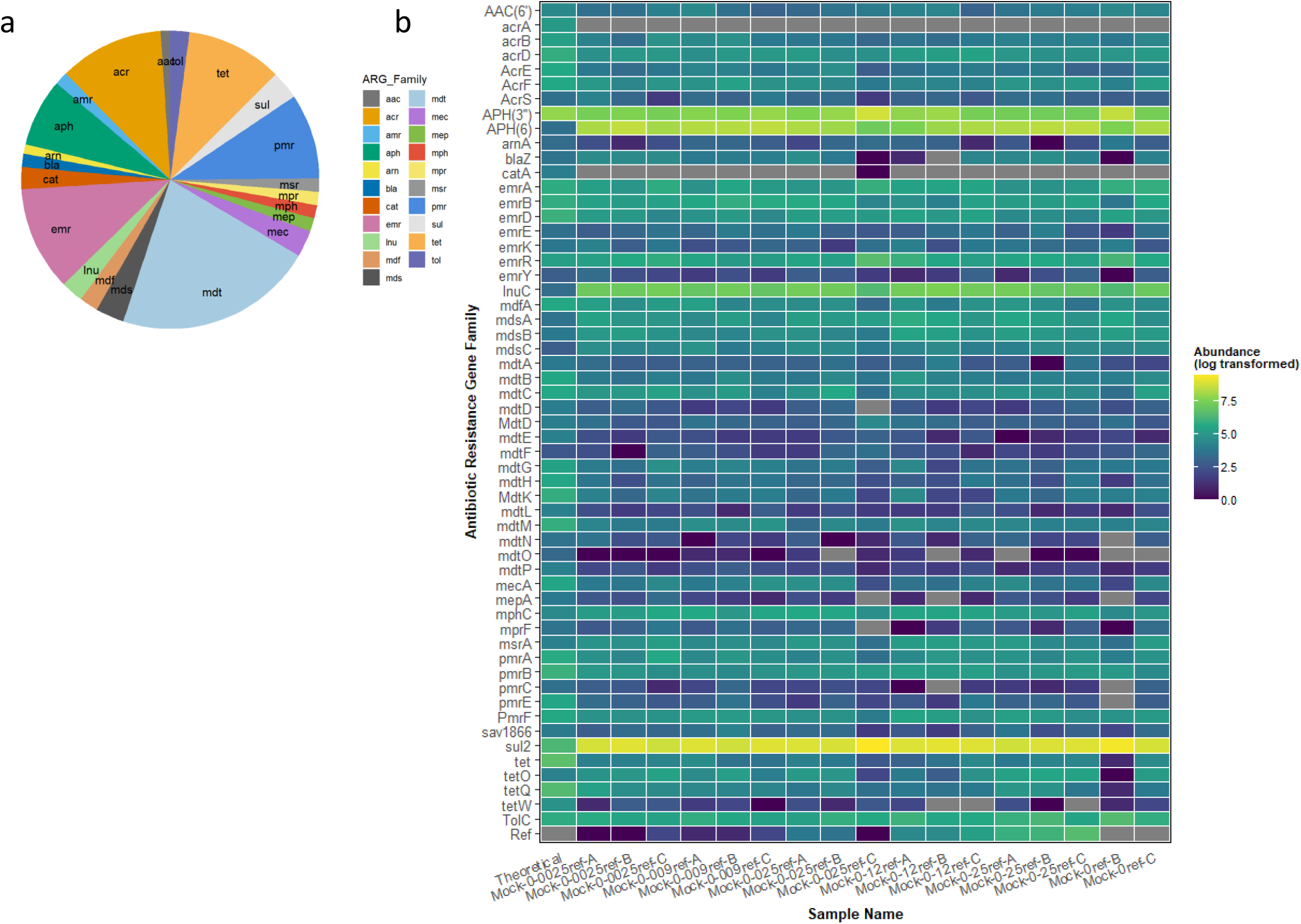
Presence and distribution of known ARGs within mock community samples. A.) Proportion of the resistome represented by each ARG Class. b.) Heatmap showing the log-transformed normalized abundance of each ARG Family from each mock sample, as well as the theoretical distribution.

### DARTE-QM differentiated resistomes between environmental sources

DARTE-QM detected 240 ARG targets across all samples in this study (including 121 in Soil-A, 172 in Soil-B, 182 in Soil-C, 202 in Swine Manure-A, 129 in Swine Manure-B, 178 in the Swine Fecal samples, and 156 in the mock-community, Supp. Table 4). Distinctions in the composition of resistomes were detected, not just from the presence of unique ARG targets but also from the abundance of the ARGs that composed the resistomes from each environment (Figure 3a). Ordination, via principal coordinate analysis based on Bray-Curtis distances of observed ARGs targets, showed clear separation of environmental sources, with the first two eigenvalues accounting for nearly 80% of the total variation (Figure 3b). Permutational multivariate analysis of variance (PERMANOVA) was used as a non-parametric multivariate statistical test to compare the variation of samples and environmental source. The results of the PERMANOVA test corroborated the apparent findings of the PCoA, and environmental sources were associated with a significant (F=11.45, R^2^=0.70, p < 0.001) portion of variation observed in the resistome profiles. DARTE-QM identified specific ARG patterns which distinguished resistomes sourced from different environmental samples, the most notable of which was within swine fecal samples where a distinctive presence of genes related to lincosamide and aminoglycoside resistance were observed. In the soils, with varied field management histories of swine and bovine manure amendment (soils B and C), we observed distinct characteristics of resistomes as well. Bovine manure-associated soils were found to be enriched with genes associated with resistance to aminoglycosides and sulfanomides, whereas the swine manure-amended soils were replete with aminoglycoside, lincosamides, and erythromycin-resistance related genes.

**Figure 3.**
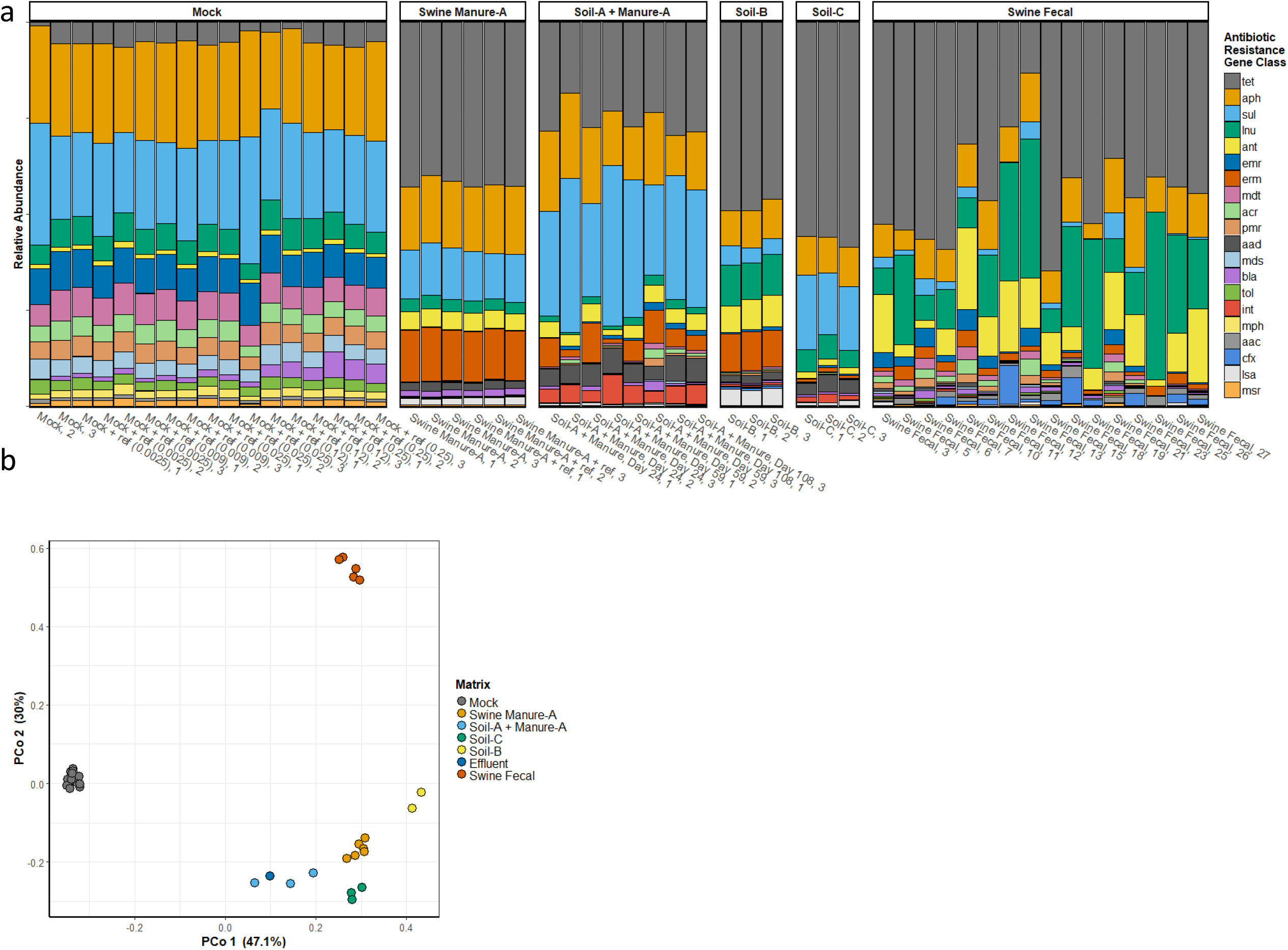
ARG profiles by source matrix. a) Relative abundance of ARG classes identified for all mock community, swine fecal, soil, swine manure, and manure-treated soils. b) Principal coordinate analysis based on Bray-Curtis distances of for resistomes, for samples passing all QC-filtering.

### DARTE-QM produced results with comparable resolution to that of metagenomes

Soil-column samples used in this study had been previously characterized through metagenome sequencing^19^ (NCBI SRA Study SRP193066). DNA from the same sources were used for sequencing with DARTE-QM study for comparison of the two methods. Metagenomes from the soil samples had an average of 241 ARG reads and were excluded from analysis; DARTE-QM returned a mean abundance of 5,839 ARG reads in those same samples. Four swine-manure samples from the metagenome study yielded a mean ARG abundance of 76,226 reads, and the 12 manure treated soil samples yielded an average of 7,377 ARG reads. DARTE-QM produced mean abundances of 32,678 and 13,488 ARG reads in the same samples.

Relative abundance of ARG classes showed similar profiles for swine-manure from both technologies. DARTE-QM reads were classified into 99 ARG families and metagenome reads to 56 ARG families. From those, 39 ARG families were shared between the two methods and accounted for 89% and 84% of metagenome and DARTE-QM ARG families, respectively (Figure 4). In the manure-treated soil samples DARTE-QM identified 99 ARG families and metagenomes 92, sharing 50 of those that accounted for 90% and 83%, respectively. For identifying diverse ARGs, DARTE-QM is disadvantaged by being a targeted method. For example, the metagenomes had a noticeable presence of genes from the AMR gene families for resistance-nodulation-cell division antibiotic efflux pump (M*ux* and *Mex* ARG Classes), which were not targeted by DARTE-QM. A direct comparison of both approaches constrains the metagenomes to those targeted by the primers of DARTE-QM (Figure 4b). In this comparison, metagenomes identified 48 ARG families in the swine manure samples and 65 in the manure-treated soils. Diversity measurements using the Shannon-Weiner Index of ARG classes showed similar values between the methods with DARTE-QM having *H* = 2.95 in swine-manure samples and *H* = 2.87 in manure-treated soil samples, while metagenomes had *H* = 2.92 in swine-manure samples and *H* = 2.84 in the manure-treated soils.

### DARTE-QM can distinguish gene variants through sequencing

Two high-abundance genes, *erm35* encoding for the macrolide-lincosamide-streptogramin and *tetM for* tetracycline resistance, were selected for variant analysis. DARTE-QM reads classified as either of these genes were clustered at 97% nucleotide identity, resulting in three clusters for *erm35* and five clusters for *tetM*. Each cluster contained a minimum of ten unique sequences. The primary *erm35* cluster contained 4,785 reads (Supp. Table 6, Supp. Figure 3a). The other two *erm35* clusters were defined by 5 to 10 base pair variations within the associated 13 and 18 reads. Similarly, from a total of 24,653 reads classified as *tetM*, 96% defined the primary cluster, which was identical to one of the 6 *tetM* primer targets. Four of the other clusters, which contained between 32 and 676 reads, were defined by 9 and 24 base pair variations (Supp. Table 6, Supp Figure 3b). Bacterial hosts associated with the observed *erm35* variants were solely associated with *Bacteroides coprosuis* and *Bacteroides spp*. and is consistent with the limited diversity of known isolates carrying this gene. In contrast, the sequences associated with *tetM* clusters are known to originate in various taxa. The largest *tetM* cluster was found to be highly conserved across a broad diversity of Gram-positive and some Gram-negative isolates. In comparison, the *tetM* cluster containing 676 reads, was primarily associated with plasmids found in *E. coli* and *Salmonella*. The lower abundance of this cluster in the DARTE-QM data is consistent with the low relative abundance of Enterobacterales in swine gut-associated samples^20^. Similarly, the other *tetM* clusters were associated with *Streptococcus* strains and found in a lower diversity of taxa compared with the largest cluster.

## DISCUSSION

DARTE-QM was conceptualized as an approach towards more efficient characterization of ARGs found in microbiomes. Specifically, we developed DARTE-QM to address the cost limitations of metagenomic approaches for ARG monitoring in environmental samples, where ARGs of interest often require significant sequencing depth and coverage. One of the major goals was to drastically scale the number of samples able to be evaluated by leveraging the high-throughput capabilities of barcode-multiplexing combined with amplicon library preparation. Similar to other amplicon-sequencing platforms, the costs of DARTE-QM are driven by the synthesis of primers and the price of sequencing. As DARTE-QM targets specified genes for amplification, it is able to enrich and detect ARGs that are present in low abundance, which is often a barrier for shotgun metagenomics. The number of samples that can be processed using DARTE-QM is limited by the number of unique barcode sequence adapters, the sequencing depth required per sample, and the number of gene targets. At the time of this study, the number of gene targets was constrained by the TruSeq platform, which currently supports 1,536 primers and 96 barcoded samples.

The aim of this study was to demonstrate the efficacy of DARTE-QM for characterizing ARGs from environmental samples. Our results showed that DARTE-QM had success detecting the presence of hundreds of diverse ARGs across soil, manure, water, and our mock-community samples. While DARTE-QM was designed with the capacity to identify diverse ARG targets, our assessment was limited by ARGs contained in our samples. We used DNA extracted from isolates with known genomes and ARG distributions to evaluate the sensitivity and accuracy of DARTE-QM. We observed strong performance for detecting ARGs in our mock-community, having 98% of ARGs detected with high sensitivity and specificity. There was evidence of DARTE-QM’s ability to quantify ARG presence with the correlation of abundance to varying concentrations of our synthetic oligonucleotide reference in the mock-community samples. Those results, though not a perfect correlation (r^2^ = 0.91), illustrate that DARTE-QM is affected more by the amount of DNA available for the primer than by the competition between primers to find targets. Finally, comparisons to metagenomes suggested that DARTE-QM could detect similar measures of diversity of ARGs from samples. While the distributions of ARGs within the resistomes varied between DARTE-QM and metagenome resistomes, the differences between environmental sources could be distinguished, and broad patterns of resistance classes were similar. Combined, these results confirm that the primers used for DARTE-QM successfully amplified ARGs despite the potential for interference when simultaneously amplifying multiple gene targets in uniform conditions.

In cases where DARTE-QM abundances varied the most from expected, the gene targets were often associated with plasmids and other mobile elements. Multiple copies of these genes may exist per cell and result in the underestimation of these genes. For instance, *aph3-ib, aph6-id* and *sul2* are found on the same IncQ plasmid. This is a likely reason for the results of much higher observed copy numbers than other ARGs, as well as the theoretical estimate. The IncQ plasmid has been reported to have anywhere between 10 to 16 copies per cell.^21^ The gene *aph(3’)-IIa*, is located on an IncI2 plasmid, which conversely is a low copy number plasmid^22^, and is consistent with our results. The optimization of future versions of this platform for specific gene targets is possible. In the case of plasmid-associated genes or genes for which amplification failed, PCR conditions could be varied for optimal amplification and specific gene standards could be included for absolute quantification. Further, it is possible to select primers for DARTE-QM to target specific resistance classes, rather than the broad array of targets demonstrated in this study.

A limitation of DARTE-QM is the presence of biased PCR amplification and associated amplicon artifacts. These sequencing artifacts were observed in all samples in this study and could be distinguished by the presence of a primer with an untargeted sequence. While these genes could be non-specific amplification of primers targeting other biological genes, the presence of poly-A and poly-T sequence patterns, like those seen in single cell amplification^23^, along with their majority singleton presence, suggested that they were sequencing artifacts. While these artifacts present an impediment for leveraging the sequencing coverage of DARTE-QM, we found that with at least 25,000 reads per sample, we could identify 90% of the ARGs present in mock-community samples. These sequencing artifacts also seemed to be produced by particular primers and in samples from specific environments, suggesting opportunities for optimization in future development of DARTE-QM. For instance, the primers targeting vancomycin-associated ARGs produced large number of artifact reads, and no vancomycin ARGs were expected in any of our samples. Similarly, many of the samples that produced the highest percentage of reads as artifacts were from soils, a medium known to have PCR inhibitors^24^. In samples where there was high-quality DNA and lower diversity (e.g., mock-community samples), it did not appear that the artifacts obstructed the production of true-positive reads. For screening of a broad range of diverse environments, artifacts are easily filtered through target alignment and classification. Future studies aimed at improving the sequencing library preparation protocols for sample types or ineffective primers will continue to improve the platform based on the knowledge gained.

The most beneficial aspect of DARTE-QM to improving microbiome ARG monitoring is its ability to detect ARGs at costs that will allow hundreds of samples to be screened simultaneously. A current challenge to antimicrobial resistance monitoring is that characterizing broad indicators are expensive, and thus it is difficult to standardize studies for monitoring. DARTE-QM is a complement to existing approaches to characterize ARGs. We envision an optimal system whereby the most relevant ARGs in a study can be detected with less bias using metagenome sequencing, and these ARGs can subsequently be targeted for numerous samples using DARTE-QM. The sequencing from DARTE-QM can then provide information on the distribution of ARGs, as well as sequence variants, in a systematic fashion, even if in low abundance.

DARTE-QM is the first demonstration of simultaneous library preparation and subsequent sequencing of hundreds of unique gene targets from environmental DNA. Here, we demonstrated this application for the characterization of ARGs and associated resistomes in environmental samples, however, DARTE-QM presents the opportunity to apply this approach towards gaining sequencing information for other diverse functional genes as well. This platform is particularly suited for studies in which genes of interest are numerous and well-defined, and where sequencing information from these genes would provide benefits to understanding biological operations (e.g., point mutations or association with sequences with host information). The ability to affordably scale for numerous genes and samples provides a much-needed resource for not only the field of antimicrobial resistance but for researchers interested in scaling functional gene characterization. Finally, we recognize that this is the first evaluation of DARTE-QM and that there are significant opportunities to further develop this approach for more targeted study. Given the simultaneous amplification of primers in DARTE-QM, we expect that the more specific the gene targets, the more optimized the library preparation can be for reliable quantification.

## DATA AVAILABILITY

Sequence files, sample metadata, and the genome sequence for the mock-community member sequenced by the USDA facility in Ames, IA, can be found through FileShare this link https://doi.org/10.25380/iastate.14390342

Alternatively, all metadata and mock genomes used it the study are available through the same repository as the code for analysis.

## CODE AVAILABILITY

All code used for processing and analysis is open-source and can be found at https://schuyler-smith.github.io/DARTE-QM/

## ONLINE METHODS

### Sequencing Targets and Primer Design

Antibiotic resistance gene (ARG) targets for primer design were chosen and aggregated from two sources. There were 2,472 sequences were obtained from the ResFinder database (version 3.2, November, 2016)^25^, associated with 67 antibiotic resistance families. ResFinder was selected on account of its manual curation of genes associated with acquired antibiotic resistance. An additional 409 ARG-associated sequences chosen as well, which had previously demonstrated high prevalence in animal agriculture^19^. To abide with the limitation of the number of allowed primers with the Illumina TruSeq library preparation, later described, the conglomerate of the chosen sequences was ultimately curated to representative sequences that targeted genes deemed of most interest to antibiotic resistance in agriculture. A single 300 bp synthetic oligonucleotide sequence was designed for use as a reference (reference target gene in Supp. Table 1). The synthetic oligonucleotide was designed with no biological context to ensure that it would not interfere with any ARG detection, save for appropriate restriction sites that were added to allow for insertion into a pUC19 cloning vector. The sequence was compared to the entirety of the NCBI Genbank database and was confirmed to share no significant similarity to any existing records. Lastly, we included 25 sequences based on those used by the Earth Microbiome Project^26^ to target the V4 variable region of the 16S rRNA gene.

The goal of primer design was to target the maximum number of our chosen sequences, with the highest specificity, staying within the set limit of 1,536 primers for the library preparation. Primers were designed using the Ribosomal Database Project’s EcoFunPrimer software:^27^ product minimum length = 220, product maximum length = 330, Oligo minimum size = 22, oligo maximum size = 30, maximum mismatch = 0, temperature minimum = 55, temperature maximum = 63, hair-pin max = 24, homo-max = 35, assaymax = 30, degenmax = 6, noTEendfileter = T, nopoly3GCfilter = T, polyrunfilter = 4, GCfilter min = 0.15 GCfilter max = 0.8. This produced 1,340 primers (Supp. Table 1) to target the ARG associated sequences, which accounted for 2,184 sequences (88.3%) from those selected (Supp. Table 2). Two primers were created for the synthetic oligonucleotide, and 30 were included for targeting all degeneracies of the 25 16S rRNA sequences. In total, DARTE-QM used 1,372 primers (668 forward-primers, 704 reverse-primers) for 796 primer pairs to be used with Illumina’s TruSeq Custom Amplicon Low Input library preparation. These primers targeted representative sequences of all 67 antibiotic resistant families and 662 ARGs.

### Library Prep

Oligonucleotide primers were created in Illumina Design Studio and ordered through Illumina (Supp. Table 1). Paired-end libraries for each sample were prepared using the TruSeq Custom Amplicon Low Input Kit (Illumina) according to the manufacturer’s instructions. This kit allows generation of up to 1536 amplicon targets over 96 samples. All DNA was diluted to 10 ng/uL during library preparation, or prepared with no dilution where concentrations were less than 10 ng/uL. An Agilent High Sensitivity D1000 ScreenTape System (Agilent Technologies) was used for measuring DNA concentration of prepared libraries. For sequencing, the MiSeq Reagent Kit v2 (300-cycles) (Illumina) reagents were used with the MiSeq sequencing platform.

### Samples

#### Mock-community

A mock-community composed of 20 cultured isolates^28^ was created for purpose of assessing the effectiveness of DARTE-QM. Nineteen of the genomes were available from the NCBI GenBank, and a single genome was sequenced at the USDA Animal Research (Ames, Iowa) (Supp. Dataset 1). The ARGs found within the genomes were annotated using ResFam and also the Comprehensive Antibiotic Resistance Database (CARD, version 2.0.1)^29^. We included 6 mock-community samples sequenced in triple replicates with 0, 0.0025, 0.009, 0.025, 0.12, 0.25 ng of the synthetic oligonucleotide reference.

To evaluate the practical implementation of DARTE-QM using environmental samples, we used 19 environmental samples originating from intrinsic and manure-amended soils, swine manures, effluent from manure-amended soils, and swine fecal samples that passed quality filters. Samples were selected from two previously published studies. In the first study, laboratory soil columns and rainfall simulations were used to evaluate the influence of swine manure amendment on soils and effluent^19^ (Supp. Table 4). In the second study, fecal samples from swine with varying antibiotic usage and routes of administration were used^30^. Samples from a subsequent laboratory soil column experiment designed to evaluate the influence of either swine or beef manure on soils and effluent were also included. Metagenomes were available for 14 samples (NCBI SRA database Bioproject PRJNA533779) were used for comparisons to DARTE-QM results.

#### Data Analysis

All analysis was done in the statistical language R, unless otherwise stated. DARTE-QM sequences were quality checked using FastQC (v0.11.9)^31^ (Figure 1). Reads were demultiplexed by primer, which were identified and removed using Cutadapt (v2.10)^32^ with an error tolerance of 0.1 and a phred-score quality threshold of 20^32^. High-quality paired-end reads were merged using PEAR (v0.9.8)^33^, requiring a minimum overlap of 10 bp. Merged reads were aligned against our database of targeted sequences using BLAST (v2.10)^34^. Successful alignment required a minimum of 90 bp and 98% similarity. For paired-end reads that were not able to be merged, each was aligned to the target-database individually. If both reads aligned to the same target, the read with the longest alignment was selected as the representative sequence. We defined a successful amplification as a read for which a primer sequence was present, and the amplified sequence aligned to the primer’s intended target with at least 90 bp length and at least 95% identity. Reads identified as having 16S rRNA primers were classified using the RDP Classifier^35^ with default parameters, and then unpaired reads selected in the same manner as for ARGs.

Each read was classified as either true positive (TP), false positive (FP), false negative (FN), or true negative (TN). This classification was based on alignment to the ARG reference database, where reads were deemed to be TP when both the intended primer target and read sequence aligned to the same gene; FP when the primer target and the read sequence did not agree; FN when no primer was found but the read sequence was able to be aligned to a reference ARG; and TN reads were assigned as all reads within a sample assigned as TP outside of the primer in question, i.e., all reads that were correctly identified as not being the targeted read. Success for DARTE-QM was evaluated on three metrics: sensitivity (TP/[TP + FN]), specificity (TN/[TN + FP]), and accuracy ([TP + TN]/[TP + FN + TN + FP]) for each gene target (i.e., primer) and each sample.

The ability of DARTE-QM to quantify ARG presence was tested by comparing observed counts of the synthetic oligonucleotide to the expected concentrations. Samples were normalized by rarefying to a sequence count of 5,000. Samples with a sequence count less than 5,000 were discarded. Alpha diversity, richness, of ARGs was calculated using Shannon’s index. Principal coordinate analysis was conducted to evaluate the variations of resistome profile in samples. Based on the relative abundance of ARGs in each sample, Bray-Curtis distances were calculated for each pair of samples, and the first two components of the eigenvalue decomposition were plotted. Permutational multivariate analysis of variance (PERMANOVA) was used to identify the significant factors (e.g., experiments, source-matrices) which contributed to the observed resistome variation. Cluster analysis was performed using k-means.

#### Variant Analysis

To evaluate the presence of gene sequence variants, the observations of variants were estimated for sequences associated with the *erm35* and *tetM* genes. The forward reads of sequences which aligned to the DARTE-QM gene targets were clustered at 97% sequence similarity with CD-HIT (v4.6.7)^36^. Clusters containing greater than ten sequences were considered in our results, with representative sequences for each cluster determined by CD-HIT. Alignment was performed and visualized with JalView using ClustalW (v2.11.1.3)^37^.

## ACKNOWLEDGEMENTS

This project was supported (or partially supported) by AFRI food safety grant no. 2016-68003-24604 from the USDA National Institute of Food and Agriculture. We thank Jennifer Jones and Kathy Mou at the *ARS-USDA National Animal Disease Center* for their help with library preparation. We thank Jared Shelerud at Illumina for his help with the TruSeq Custom Amplicon platform.

## AUTHOR CONTRIBUTIONS

A.H., H.A., M.S., F.Y., N.R., and J.C. were designed the project; A.H. H.A., M.S., and S.H.-L. were involved in funding-acquisition; S.S. analyzed the data and wrote the manuscript with assistance from A.H. and N.R.

## ADDITIONAL INFORMATION

Supplemental information

## PEER REVIEW INFORMATION

**Supp Figure 1.**
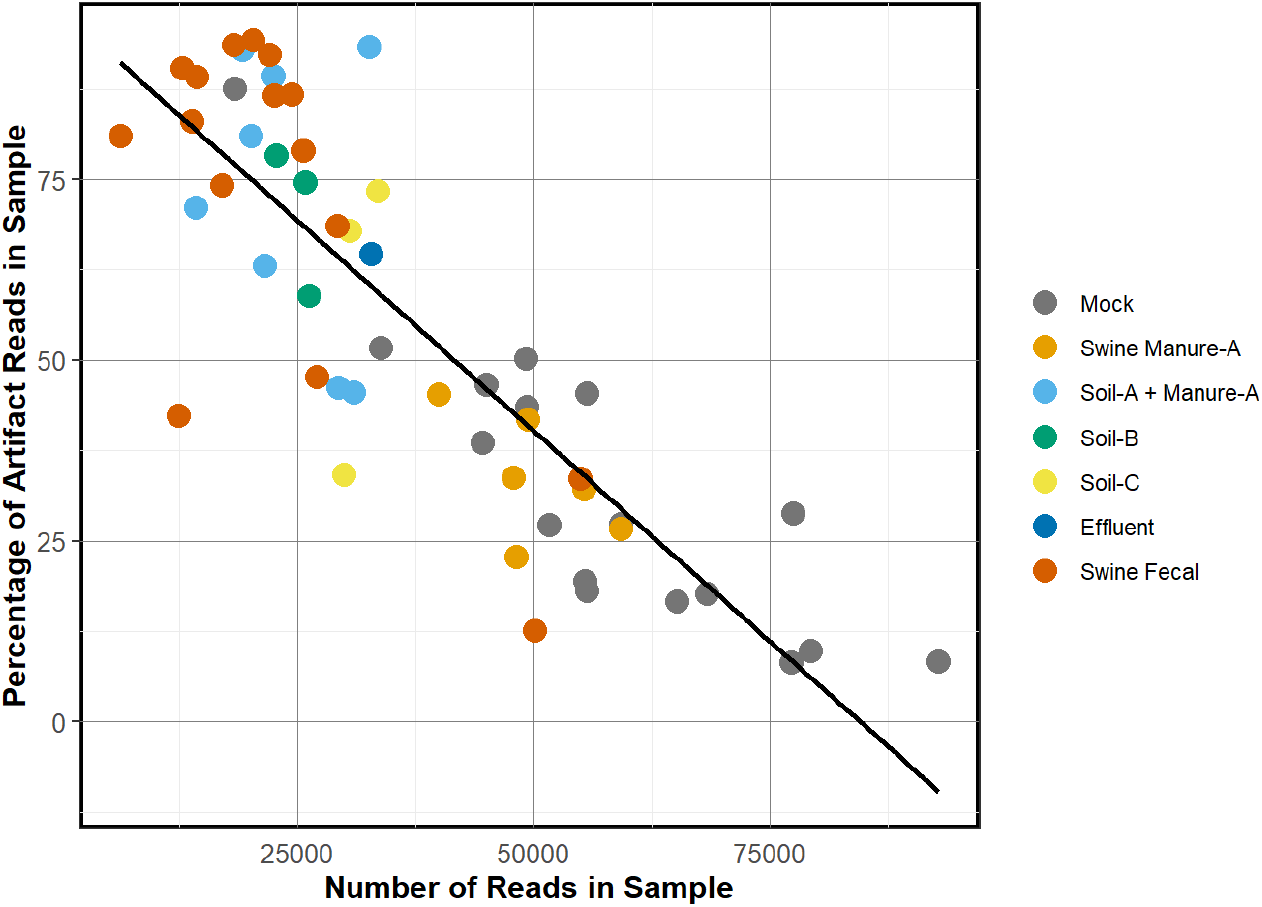
Linear correlation of the percentage of artifacts reads present in a sample to the total number of reads in the sample. Reads were defined as sequencing artifacts if a primer was located on the 5’ end of the sequences and the read did not align to any of reference ARGs or any other location in the mock-community genomes. The percentage of sequencing artifacts observed was higher for environmental samples relative to mock community samples and was also inversely correlated (R ^2^= 0.68) to the number of reads in a sample.

**Supp Figure 2.**
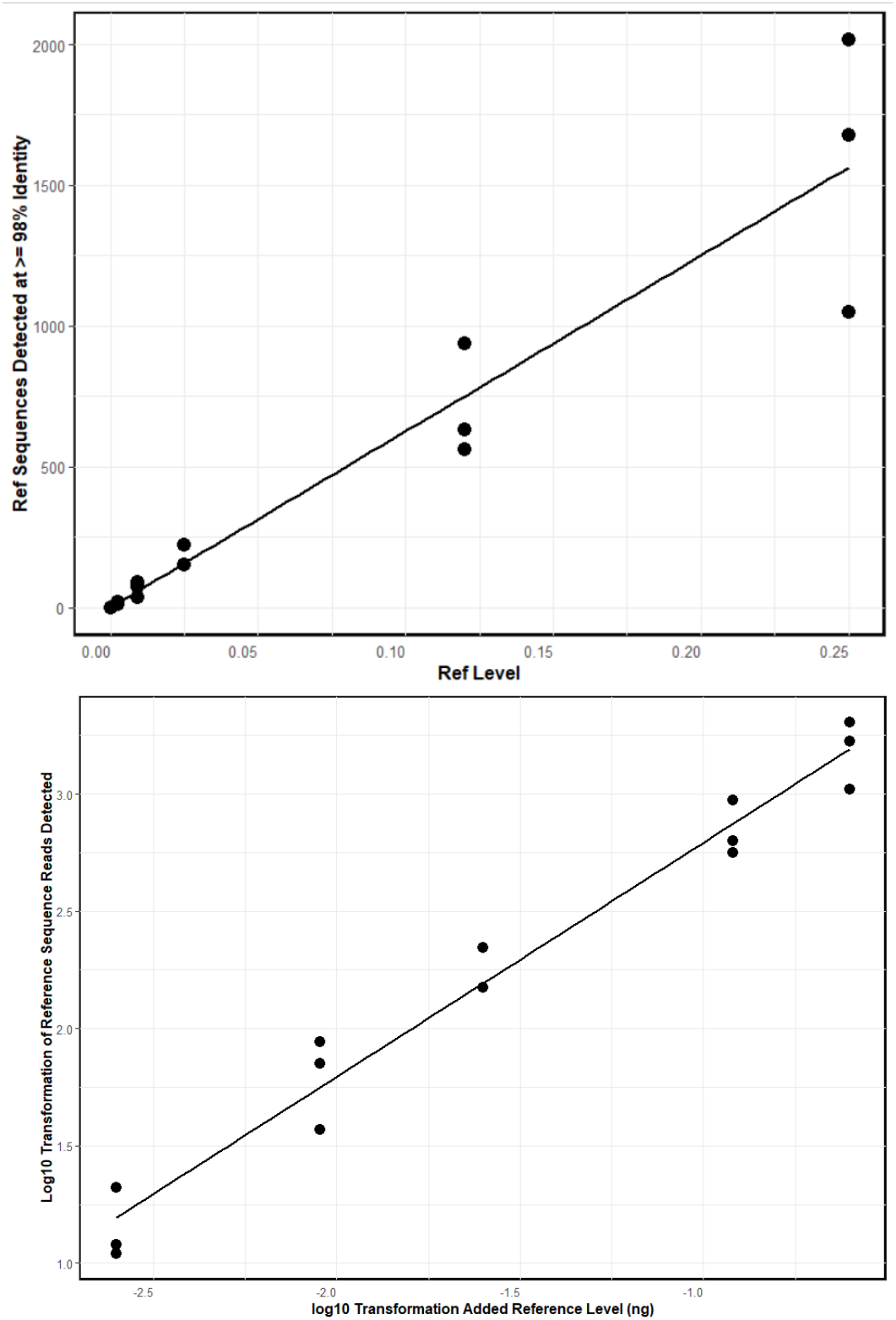
Linear correlation between the concentration of the reference sequence added to mock community samples and the number of reads which aligned to the reference sequence. The linear model found there there to be a strong correlation (R^2^ = 0.91), indicating DARTE-QM iwas sensitive to DNA quantity.

**Supp Figure 3.**
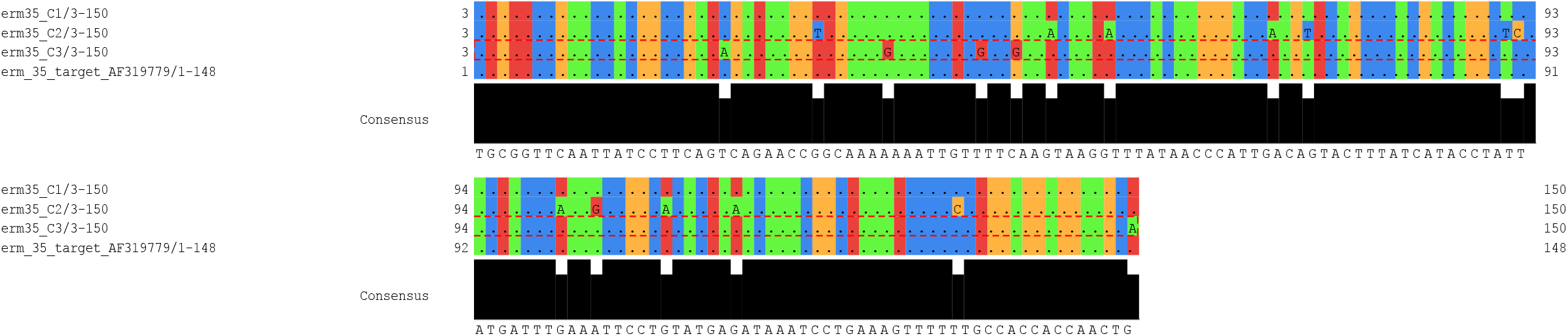
a) Alignment of gene targets and sequences identified by DARTE-QM gene target for a) *erm35* and [continued on next slide]

**Supp Figure 3.**
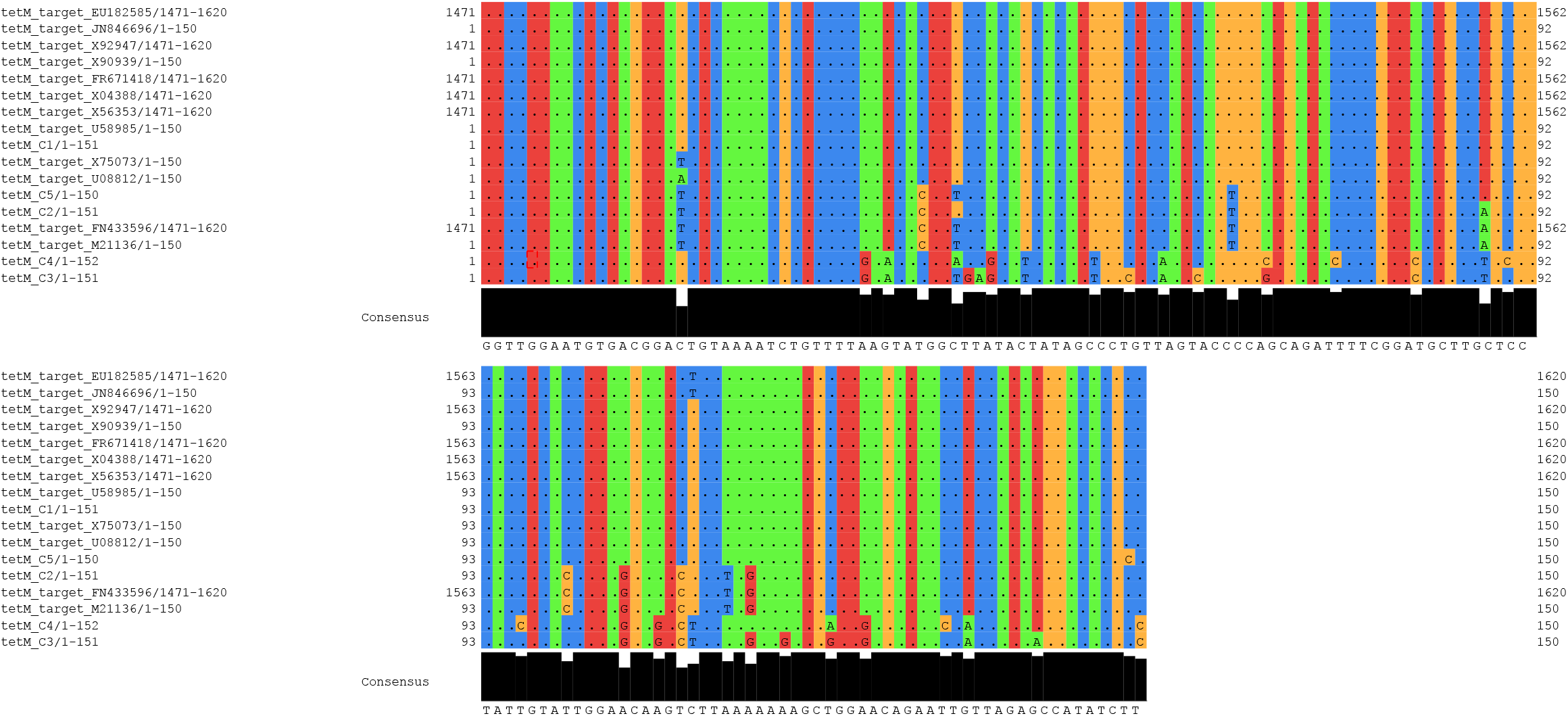
b*) tetM* genes. Sequences are representative sequences identified for clusters of reads by 97% sequence identity (see also Supp. Table 7). The presence of gene variants are shown in the corresponding three clusters for *erm35* (erm35*_C1-C3)* and five clusters for *tetM* (*tetM_C1-C5)*. Genes targeted by DARTE-QM are also shown.

