## Supplementary material for "Diversity of Antibiotic Resistance genes and Transfer Elements-Quantitative Monitoring (DARTE-QM): a method for detection of antimicrobial resistance in environmental samples": Figure Captions

Supp. Table 1. Primer design information for the 798 primer pairs created for evaluating DARTE-QM including the target gene (ARG class described in Supp. Table 2), the forward and reverse primer pairs, the targeted product length (bp), primer lengths (bp), primer GC percentages, and the target sequence.

Supp. Table 2. Classification of antibiotic resistance class, family and gene for each of the sequence targets for the primers of DARTE-QM.

Supp. Table 3. Genome and taxonomic information for the 20 members of the mock-community.

Supp. Table 4. Metadata for each of the samples used to evaluate DARTE-QM. The ID of the sample for DARTE-QM; the sample name is an abbreviated descriptor of the sample; the description is a brief description of the sample; the matrix is the environment of origin of the sample; the reference added is the concentration of synthetic oligonucleotide reference added to a sample; the reference is the associated publication describing a sample; passed QC denotes which samples fulfilled the criteria to be used for analysis within the study; and the total ARGs detected by DARTE-QM.

Supp. Table 5. Reads produced by DARTE-QM for each sample. Raw reads are the total number of reads produced; merged reads are the total number of merged paired-end sequences; ARG reads are the number sequences annotated to a reference ARG; unique ARGs are the total number of unique ARG sequences; ARGs are the total unique types of ARGs observed per sample; and 16S SSU are the total number of reads associated with 16S rRNA SSSU primers.

Supp. Table 6. Classifications of reads produced from each primer pair in each sample based on metrics of success, including true positives (true_pos), false positives (false_pos), false negative false_neg), and true negatives (tru_neg). The total number of reads (n_reads) and artifactual reads are shown, and only reads that were not identified as artifacts were further classified.

Supp. Table 7. Gene variants for *erm35* and *tetM* identified with DARTE-QM. Gene clusters were created by clustering reads associated with these ARGs at 97% nucleotide identity, requiring a minimum of 10 observed reads. The number of base pair variations, % identity, and % coverage of gene length from the most similar alignment in NCBI was estimated.

Supp. Dataset 1. Sequenced genomes of the 20 mock-community members in FASTA format.
